# A preferred curvature-based continuum mechanics framework for modeling embryogenesis with application to *Drosophila* mesoderm invagination

**DOI:** 10.1101/029355

**Authors:** K. Khairy, W. Lemon, F. Amat, P. J. Keller

## Abstract

Mechanics plays a key role in the development of higher organisms. However, working towards an understanding of this relationship is complicated by the fact that it has proven difficult to model the link between local forces generated at the subcellular level, and tissue deformation at the whole-embryo level. Here we propose an approach first developed for lipid bilayers and cell membranes, in which force-generation at the cytoskeletal level only enters the shape mechanics calculation in the form of local changes in preferred tissue curvature. This allows us to formulate the continuum mechanics problem purely in terms of tissue strains. Relaxing the system by lowering its mechanical energy yields global morphogenetic predictions that accommodate the tendency towards this local preferred curvature, without explicitly modeling force-generating mechanisms at the molecular level. Our computational framework, which we call SPHARMMECH, extends a three-dimensional spherical harmonics parameterization known as SPHARM to combine this level of abstraction with a sparse shape representation. The integration of these two principles allows computer simulations to be performed in three dimensions, on highly complex shapes, gene expression patterns, and mechanical constraints.

We demonstrate our approach by modeling mesoderm invagination in the fruit-fly embryo, where local forces generated by the acto-myosin meshwork in the region of the future mesoderm lead to formation of a ventral tissue fold. The process is accompanied by substantial changes in cell shape and long-range cell movements. Applying SPHARM-MECH to whole-embryo live imaging data acquired with light-sheet microscopy reveals significant correlation between calculated and observed tissue movements. Our analysis predicts the observed cell shape anisotropy on the ventral side of the embryo and suggests an active mechanical role of mesoderm invagination in supporting the onset of germ-band extension.

## Introduction

Embryonic development is accompanied by fundamental changes in shape, including long-range tissue reorganization and epithelial folding. At the cellular and molecular levels, these morphogenetic processes are driven by cytoskeletal reorganization, leading to force generation (1-6). This activity is orchestrated at the whole-embryo level through differential expression of genes. In addition, mechanical constraints also determine and limit morphological configurations available to a developing organism, and the forces generated in turn influence gene expression itself (7-9). It is thought that forces are transmitted over long ranges through the material of a tissue. However, details of how forces exercise such long-range effect can only be understood by combining experimental data and computational models. With advances in microscopy technology, detailed live imaging data of whole-embryo morphogenesis have become available (10–12) (**Fig. 1a** and **Supplementary Videos 1** and **2**). Large-scale approaches to imaging and computational image analysis have produced near-comprehensive quantitative gene expression distribution maps for the early embryo (13, 14) (**Fig. 1b**). Therefore, it should be possible to predict observed tissue movements and cell shape changes that result from local force generation, by considering whole-embryo tissue mechanics.

**Figure 1.**
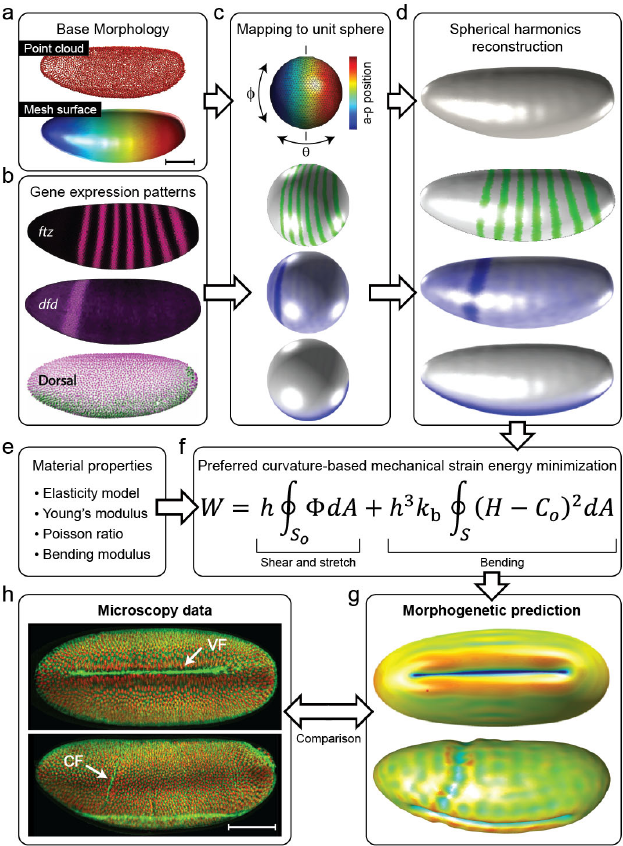
SPHARM-MECH overview. (**a**) Morphology data. Top: Cell centers from segmentation of the nuclear label His2Av-RFP channel from a three-dimensional stack of a live adaptive SiMView light-sheet microscopy recording of a whole *D. melanogaster* embryo at the cellularized blastoderm stage. Bottom: surface triangular mesh based on cell centers. Color code anterior-posterior (y-axis) with colorbar as in (c)-top. (**b**) Gene expression data. Example gene expression patterns of *ftz, dfd* and *dl. ftz* and *dfd* patterns are displayed using PointCloudXplore(13). (**c**) Spherical mapping. Cartesian coordinates (top: color code corresponds to (a) bottom with values provided in colorbar). Gene expression patterns of *ftz, dfd* and *dl* are mapped onto the unit sphere. (**d**) Spherical harmonics reconstruction (projection). Based on the spherical mapping in (c) Fourier coefficients are calculated for morphology and gene expression patterns to produce the internal analytical representation used by SPHARM-MECH (Online Methods). The outline is represented by 25 numbers (SPHARM coefficients). (**e**) Material properties data may be obtained from experimental measurements or estimated based on physical assumptions. (**f**) Energy minimization. Gene product activity increases the configuration-dependent strain energy by inducing local changes to preferred curvature. Morphology changes occur to relieve that energy (Online Methods and **Supplementary Note 6**). (**g**) Example SPHARM-MECH simulation results showing formation of the ventral furrow invagination (top) and simultaneous simulation of ventral and cephalic furrows (bottom). (**h**) Ventral and lateral views of maximum intensity projections of three dimensional stacks of an adaptive SiMView microscopy recording of a whole *D. melanogaster* embryo, homozygous for the membrane label Spider-GFP and the nuclear label His2Av-RFP, 22 min after onset of gastrulation (see **Supplementary Videos 1** and **2** for complete time-lapse recording). Scale bars, 100 μm (a, h).

There have been a number of efforts to model morphogenesis on the computer (15); for example: in the fruit fly (16–21), the sea urchin (22, 23), and the chick embryo (24, 25). However, accurate three-dimensional modeling of embryogenesis remains a difficult task. A major challenge is the requirement of a multi-dimensional mathematical representation that is able to express embryo shape, gene expression patterns and material properties accurately, and in a manner that allows efficient numerical application of constitutive tissue behavior. A second challenge is to link force generating mechanisms, which occur at the subcellular level through cytoskeletal dynamics, to a continuum mechanics formulation, which is necessary for performing whole-embryo simulations in realistic computation times. This link should provide a level of abstraction that is able to approximate the activities of various cytoskeleton-level mechanisms, and yet communicates their effects on a macroscopic scale in a fashion compatible with standard mechanics treatments.

To address the above challenges, we developed an open-source computational biomechanics framework which we call SPHARM-MECH (**Fig. 1**). Shape representation is based on a powerful Fourier basis for approximating functions on the sphere known as the spherical harmonics basis functions (**Supplementary Fig. 1** and **Supplementary Note 1**). The spherical harmonics have been used to represent, register and compare complex morphologies of simply-connected objects through an approach known as SPHARM (26–28). SPHARM-MECH data representation generalizes the SPHARM approach. It concisely encodes, in addition to morphology, gene expression distributions to facilitate data-driven identification of regions of local gene-product activity (**Fig. 1a–d** and **Methods**). Its fidelity to experimental data is limited only by the spatial and temporal resolution of the recording device and noise. Harmonic functions of higher order gradually provide more detail, as expected from a converging Fourier series (**Supplementary Fig. 2**). Importantly, gene expression patterns are naturally mapped onto morphologies independent of their source in a straight-forward manner (**Fig. 1b–d, Supplementary Fig. 3a–g** and **Supplementary Note 5**). Using these maps, regions of local mechanical activity are generated (**Fig. 2a** and **Supplementary Fig. 3h**).

**Figure 2.**
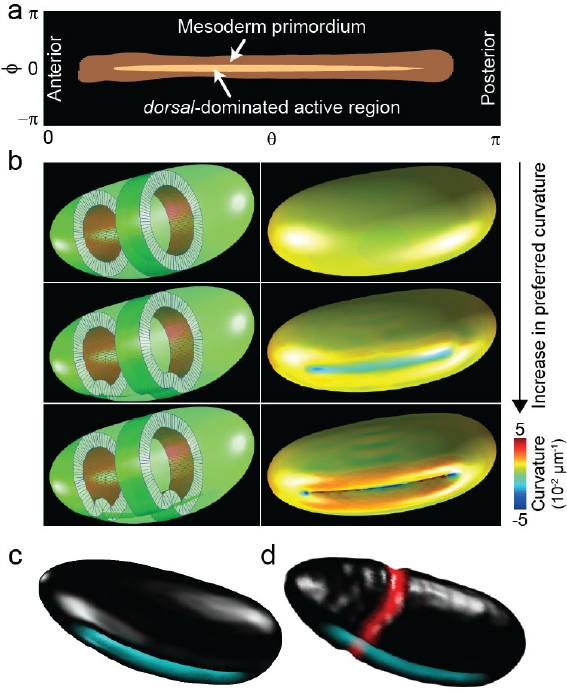
Simulation of ventral and cephalic furrows. (**a**) Mesoderm primordium region: Threshold of map that results from the interaction of *snail, twist* and *huckebein.* Threshold was chosen so the mesoderm primordium covers 16% of the total area. Area in the center shows the 8-cell wide region defined as the activity map, based on *dorsal* and *huckebein* expression patterns, used for VFI simulations in (b and c) (**Supplementary Note 8**). Angles *θ* and *ϕ* represent the anterior-posterior and dorso-ventral positions respectively. (**b**) Left: Perspective views of VFI morphology predicted by SPHARM-MECH when gradually increasing the preferred curvature. Right: local mean curvature color code of corresponding morphologies on the left. (**c**) Perspective view of the bottom-most morphology of (b) with color showing the active region used. (**d**) Simultaneous simulation of VFI and CF. Colors mark the active regions.

**Figure 3.**
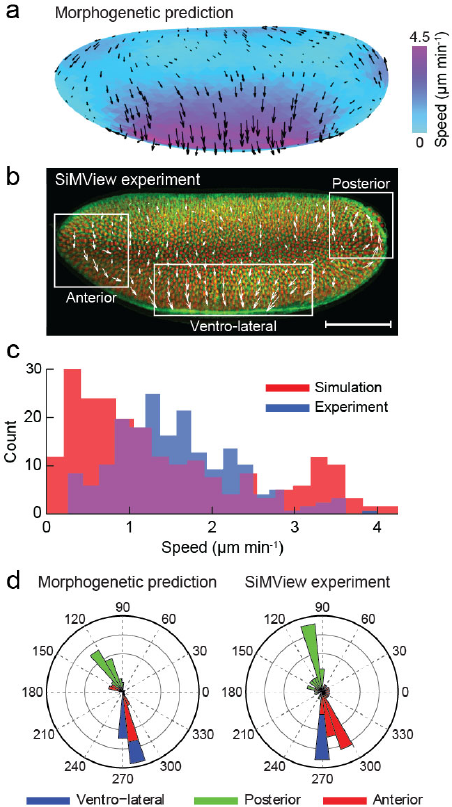
Comparison of experimentally observed and simulated tissue flows. (**a**) Tissue flow velocity field obtained from a VFI-only simulation showing speeds (color code and relative arrow size) and direction of flow. Results represent a simulation from undeformed geometry to one that equals the furrow after 10 minutes of onset of gastrulation. (**b**) Experimentally observed tissue flow velocity field obtained by manual tracking of 270 cell centroids and interpolated over the image over 10 minutes of onset of gastrulation. (**c**) Speed histogram corresponding to simulation and experiment. (**d**) Flow direction histogram angular plots corresponding to simulation (left) and experiment (right), showing resulting flow directions in the three regions outlined in (b). Scale bar, 100 μm.

Given gene activity maps, a starting morphology (**Fig. 1d**), and material properties (**Fig. 1e**), SPHARM-MECH calculates a configuration-dependent strain energy for the whole tissue (**Fig. 1f** and **Methods**). We propose that the driving force for tissue invagination and epithelial folds manifests itself mechanically as a departure of the preferred tissue curvature from its current curvature at locations of gene product activity. This can occur via a number of different molecular and cellular mechanisms, the details of which do not enter our analysis. Examples would include contraction of an acto-myosin meshwork driving the ventral furrow invagination (VFI) (**Fig. 1h**) via apical cell constriction (29), or repositioning of adherence junctions responsible for formation of a dorsal fold (30), both observed in the fruit-fly embryo. Abstraction/Omission of these details is intentional in our approach in favor of computational feasibility of whole embryo mechanics modeling. The preferred curvature concept provides the necessary link between effects of local biological force-generating mechanism and a continuum tissue mechanics formulation. When forces are generated locally, the organism is in a high mechanical energy state. Morphological changes occur in order to relieve this strained tissue configuration (**Fig. 1f**). SPHARM-MECH predicts a morphology that minimizes mechanical energy (**Fig. 1g**) by using standard methods of numerical optimization (**Methods**).

To demonstrate our approach we focus on the fruit-fly embryo. We apply SPHARM-MECH to model VFI formation in *Drosophila.* In this system, the egg shell and vitelline membrane act as additional mechanical constraints. We explicitly include the vitelline membrane as a hard mechanical constraint surrounding the embryo.

We show that forces generated by the isotropic contraction of the acto-myosin meshwork local to the future mesoderm, are responsible for observed tissue anisotropy on the ventral side of the embryo. We also show that these forces might encourage tissue movements of the posterior pole in the dorsal direction, thereby supporting the first phase of germ-band extension, and that this effect is directly related to the geometry of the egg.

## Methods

### The Software

SPHARM-MECH has been designed to enable efficient prediction of morphogenetic changes at the whole-embryo scale. This computational framework expresses the shape outline (**Fig. 1a**) and gene expression patterns (**Fig. 1b**) in terms of a small number of shape descriptors using the spherical harmonics basis functions (**Fig. 1d**), calculates a configurational strain energy based on continuum shell mechanics (**Fig. 1f**), and yields a predicted configuration that minimizes this energy (**Fig. 1g**) by using numerical optimization. SPHARM-MECH solves the mechanical problem purely in terms of strains, which is desirable, because strains constitute a readily available observable in microscopy experiments.

The central component of the software is a “shape_tools” library, that defines C++ classes for (a) a spherical triangular mesh, which is used for fast display, surface intersection tests and approximate shape properties, (b) a spherical harmonics basis, which provides accurate and efficient values of basis functions as well as first and second derivatives defined on a Gaussian quadrature grid, (c) a spherical harmonics surface class, provides accurate and efficient shape properties calculations, and (d) a shell class for continuum mechanics calculations. We also provide a development Matlab library with similar functionality.

Two applications with a graphical user interface are provided: (1) SHAPE (Spherical Harmonics Parameterization Explorer) is a utility for inspecting, modifying and exporting shapes, and (2) SPHARM-MECH, providing an interface that facilitates importing constraint and starting shapes, configuring material parameters, defining sites of mechanical activity based on gene expression, and executing energy minimization.

Details about shape properties and mechanical energy calculations are provided below (and additional details in **Supplementary Notes 1-6**).

### SPHARM-MECH spatial scheme

Embryo morphology and gene expression pattern are represented parametrically in functional form as

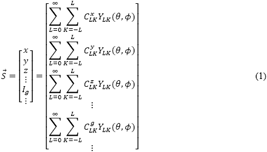

where *θ* and *ϕ* are spherical polar coordinates on an abstract unit sphere. *Y_LK_*(*θ,ϕ*)*s* are spherical harmonics functions of order *L* and degree *K* (**Supplementary Note 1**). The surface outline (in form of Cartesian coordinates) is encoded in coefficients 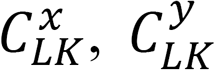 and 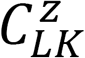. Scalar fields *I_g_*, encoded in coefficients 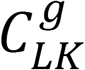, represent gene expression pattern intensities, for example: *I_snl_*, *I_twi_*, *I_hkb_* and *I_dfd_*, as well as the activity maps that determine location and extent of changes in preferred curvature from that of an undeformed morphology. In principle, SPHARM-MECH is not limited to gene expression or activity maps, but may include other field characteristics, for example position-dependent material properties.

The total number of coefficients for a shape outline is 3 × (*L*_max_ +1)^2^, where *L*_max_ is the maximum order of the series. The infinite sum in Eq. 1 is truncated at *L*_max_ = 36 in all simulations shown in this work. This upper limit provides sufficient accuracy, even for patterns that are highly localized, such as *ftz* and *eve* (**Fig. 1d, Supplementary Fig. 3c**), and for more complex embryo morphologies (**Supplementary Fig. 2**). Optionally, mirror symmetry can be imposed to decrease the number of shape coefficients by about 40%. Total volume, local mean curvature, shear and stretch deformation are evaluated using numerically stable recursion relations (31). For details see **Supplementary Notes 2-4**.

SPHARM-MECH combines the following four characteristics: (1) A highly accurate evaluation of the spherical harmonics functions and their derivatives (31) provides reliable deformation gradient tensors and therefore shear and stretch energies, as well as accurate local mean curvatures and therefore accurate bending energies (**Supplementary Notes 2** and **3**), (2) fast evaluation of surface integrals is possible by Gaussian quadrature (**Supplementary Note 4**), (3) a concise shape representation makes it practical to use standard techniques of numerical optimization, and the descriptors have physical meaning (**Fig. 4a**), and (4) the structure of the spherical harmonics representation naturally takes advantage of symmetry (**Supplementary Note 7**). The main limitation of SPHARM-MECH is the restriction to shapes that are topologically equivalent to the sphere, i.e. of genus zero, and its global support (i.e. if only local effects and local effectors are involved, then a possibly prohibitively large number of harmonic functions would be needed for accurate calculations).

**Figure 4.**
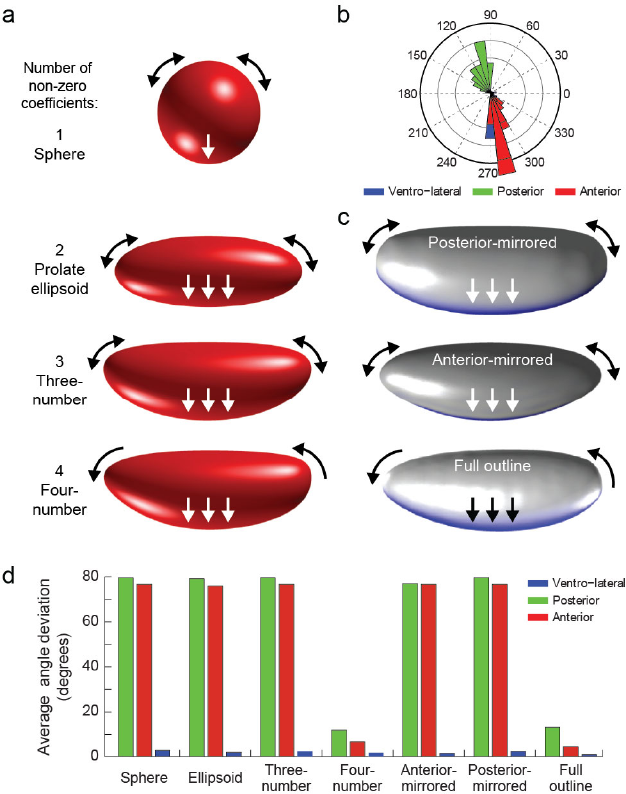
Long-range effects of embryo symmetry on tissue flows resulting from ventral furrow invagination. (**a**) Four SPHARM-MECH coefficients are sufficient to approximate the *D. melanogaster* blastoderm outline. Each coefficient breaks a different type of symmetry as seen when going from top (sphere) to bottom (four-coefficient shape). Please see **Supplementary Note 7** for more details regarding the meaning of the coefficients. When using the top three shapes (that possess left-right [anterior-posterior] symmetry) as base shapes for performing the VFI simulation instead of the full outline, tissue regions on the posterior (defined arbitrarily to be located on the right) flow dorsally only 50% of the time and ventrally otherwise. This is indicated by the black two-headed arrows, showing that the flow can go in either direction. In addition, the anterior and posterior flows are always with opposite destinations, i.e. when posterior tissue flows dorsally, anterior pole tissue flows ventrally and vice versa. However, when the anterior-posterior symmetry is broken (the four-number shape), the posterior (lower curvature) pole region tissue flows dorsally in all of the simulations (*n* = 6). This is indicated by the black single-headed arrows. Flow ventrally (towards the forming invagination) from the lateral sides is present in all cases. This is shown by the white arrows. (**b**) Flow direction histogram angular plot corresponding to a typical simulation with the sphere as base undeformed shape. Assuming the pole where tissue flows dorsally to be posterior, this shows flow directions similar to the ones observed in the experiment (and simulation with the full outline) (**Fig. 3d**). Similar results are obtained for all simulations. (**c**) Two cases were constructed: a left-right symmetric outline (top) with the posterior features preserved (posterior-mirrored) and another (middle) with the anterior features preserved (anterior-mirrored). Similar to the left-right symmetry cases in (a), the flow was observed in each direction 50% of the time. However, when performing the VFI simulation on the full morphology repeatedly, the flow was always from the posterior dorsally, indicating that it is indeed the break in left-right symmetry that biases the direction of flow. (**d**) Plot of average (*n* = 6) flow angle deviation with respect to the experimentally observed flow. Only the simulations with shapes that possess the left-right asymmetry yield differential tissue flows close to the experimental observation.

Finally, it should be noted that the spherical harmonics parameterization is not the only shape description appropriate for this kind of problem. Any complete set of orthogonal eigenfunctions defined on the sphere could be used. For highly localized effects, a polygonal mesh might be more appropriate. However, the four characteristics mentioned above provide the SPHARM-MECH framework with its particular computational utility for general embryogenesis problems.

### Mechanical strain energy

To model tissue mechanics, we use a generalized neo-Hookean constitutive material model based on the work of L.R.G. Treloar for rubber-like materials (32). The strain energy density is given by:

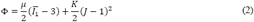

where *μ* is the second Lame constant (shear modulus), *K* the bulk modulus, *J* = *λ*_1_*λ*_2_*λ*_3_ (determinant of the left Cauchy-Green deformation tensor), and *Ῑ*_1_ (an invariant of the left Cauchy deformation tensor) given by 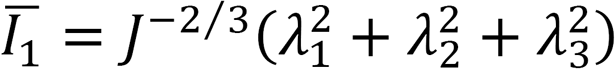. The principal stretches λ_i_ (*i* = 1, 2, 3) are calculated from shape coefficients of both current and undeformed (starting) surfaces using differential geometry (**Supplementary Notes 3** and **6**). The total contribution of shear and stretch energies is given by:

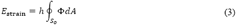

where *h* is the thickness of the shell (assumed constant) and the integration is performed over the undeformed surface *S*_o_. For information on evaluating Eq. 3 numerically see **Supplementary Note 6.** The energy contribution from bending and twisting is given by:

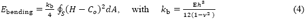

where E is Young’s modulus, ν Poisson’s ratio, *H* local mean curvature of the current configuration (**Supplementary Note 3**), and *C*_o_ the preferred local mean curvature induced by gene activity. The total energy of the shell is given by

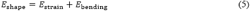

### Energy minimization

Equation 5 comprises a nonlinear system. Predicted shapes are those of minimum *E_shape_*, as found by using a constrained nonlinear function optimization SubPlex algorithm of the NLopt C++ library (33, 34). (We have included additional (optional) nonlinear optimization approaches in the SPHARM-MECH software on an experimental basis, also with the potential that other mechanical systems might require them.)

### Application example: Fruit-fly ventral furrow invagination mechanics

To demonstrate the utility of SPHARM-MECH, we applied it to the problem of whole-embryo modeling of ventral furrow invagination in *Drosophila melanogaster;* a system where significant advancements in understanding gastrulation (35–37), the associated gene expression patterns and gene product activity (13, 14) have been made.

VFI formation is accompanied by a sequence of cell shape changes that ultimately lead to mesoderm internalization (36–39). The first change in shapes of cells forming the VFI is apical constriction, which is driven by acto-myosin-contraction (29, 40). Apical constriction is local to expression of *twist* and *snail*, both of which are essential for furrow formation and are expressed ventrally along the anterior-posterior axis. This contraction is compatible with our assumption that preferred curvature is induced locally. *huckebein* sets the limits of *snail* and *twist* expression levels at the poles and *dorsal* modulates relative advancement of cells in their fate (**Supplementary Fig. 3g,h**). We calculate two different acto-myosin contractility maps (**Fig. 2a**), and perform a quasi-static mechanics simulation using each.

Gene expression patterns are calculated based on data from the VirtualEmbryo project (13), and from light-sheet microscopy recordings in case of the *dorsal* expression pattern, and are assumed constant over the simulation time-frame spanning about 10 minutes after onset of gastrulation.

### Model parameters and assumptions

Our primary model assumptions are:

a. Furrow forming forces affect the geometry through imposing a preferred curvature that is different from the preferred curvature before gene product activity starts (e.g. through contraction of cytoskeletal elements).
b. Regions of constriction cause local stiffening (larger bending resistance) relative to the passive surrounding tissue. In the chick embryo, the bending stiffness within the region of invagination-initiating cells is about twice as high as the surrounding tissue (24).
c. All tissue deforms passively according to a hyperelastic material model (Eq.2).
d. The yolk is a fluid that preserves the internal volume enclosed by the embryo during deformation and its viscosity is ignored (constant volume is enforced).
e. The vitteline membrane forms a hard constraint surrounding the embryo tissue.
f. Thin shell mechanics are sufficient to approximate the deforming surface.
g. The shell is of constant thickness *h*, i.e. *λ*_3_ = 1.
h. The shell material is nearly incompressible.
i. Non-self-intersection and non-intersection with the vitteline membrane are enforced (using a triangle-triangle intersection test (41)).

The material parameters are *ν* = 0.45 (close to that expected for water-filled tissue), *E* = 100 Pascal, *h* = 0.5 μm, *γ*= 20 for VFI and 10 for the cephalic furrow, and ratio of bending stiffness of the mesoderm primordium to that of the passive tissue *λ* = 20 for VFI and 10 for cephalic furrow.

### Results

Time-lapse, three-dimensional image data of entire *Drosophila* embryos are recorded during gastrulation, using adaptive multi-view light-sheet microscopy (42, 43). Through image processing (**Application-Specific Methods**) this provides data of cell nuclei positions and cell shape dynamics throughout this process (**Supplementary Videos 1** and **2**). This data is used to obtain a three-dimensional starting morphology driving the model, as well as the time evolution of the embryo shape outline which allows us to quantitatively assess the predictive power of the SPHARM-MECH modeling framework.

A starting surface is obtained from the outline of the blastoderm prior to the onset of gastrulation (**Fig. 2b** top). The local preferred curvature remains equal to that of the blastoderm everywhere except in regions of activity as determined by the activity map. Values of local preferred curvature within the activity region are input as multiples (γ) of the image-data-determined curvature about 10 minutes into gastrulation. In those regions, γ is increased in two steps (to 2 and 20, respectively). In each step a full numerical optimization of the strain energy is performed to yield the predicted morphology and strain field.

From previous work using electron microscopy, it is known that a core band of cells about 8-10 cell-width contracts prior to the rest of the MP (36). Using expression patterns for *dorsal* and *huckebein* (**Fig. 1b–d, Supplementary Fig. 3g,h** and **Supplementary Note 8**), we find that a map corresponding to an eight-cell wide ventral anterior-posterior oriented band (**Fig. 2a**) reproduced the initial stage of VFI qualitatively (**Fig. 2b,c**) when comparing to experimental data obtained with SiMView microscopy (**Supplementary Video 1**) and other imaging modalities (**Supplementary Fig. 4**). This is in contrast to the case when considering the activity of *snail, twist* and *huckebein* (**Supplementary Note 8**) to generate a map that is thresholded to cover 1/6 of the blastoderm surface, which is equivalent to the full mesoderm primordium region (MP) (**Fig. 2a** and **Supplementary Fig. 3h**, second from bottom). When the full MP map is used to define the region of contractility, i.e. the full MP region is contracting simultaneously, we find that the simulation does not reproduce the observed deformation even at γ = 100, but produces a frustrated ventrally slightly flattened morphology that does not invaginate. It is conceivable that constriction occurs in an out-ward fashion in which cells are activated as a function of stress induced by neighbors. We obtain the same result above when simulating cephalic and ventral furrow formation simultaneously (**Fig. 2d** and **Supplementary Note 11**).

To observe long-range effects, we perform VFI-only simulations using SPHARM-MECH (as above with γ = 20) and estimate a predicted tissue velocity field from time points zero to ~10 minutes into gastrulation (**Fig. 3a**). For comparison, experimentally observed tissue flows are extracted, by image processing, from *in vivo* whole-embryo data of morphogenesis recorded with SiMView microscopy, and covering the same time period (**Fig. 3b**). Both speeds and flow directions for simulation and experiment are compared in the form of histograms. We find very good qualitative and quantitative correspondence in material velocity fields across the entire embryo, including lateral-anterior, ventro-lateral and posterior regions that exhibit the most striking directed tissue flows (**Fig. 3c,d**). Importantly, we note the dorsal flow of tissue in the posterior pole region. This flow may appear counterintuitive at first because the simulation considered deformation resulting from VFI driving forces only, and these driving forces relate to a spatial location that is distant from the affected posterior pole region. The simulation suggests that the process that forms the ventral furrow favors a dorsal movement of posterior tissue, independent of mechanisms of germ-band extension. This result is consistent with recent findings that explicitly propose that cell shape change observed during germ-band extension is a passive response to mechanical forces caused by the invaginating mesoderm (44).

To investigate the physical origin of this differential tissue flow, we take advantage of the conciseness of the spherical harmonics shape representation. In particular, that the fly syncytial blastoderm morphology can be reasonably approximated by only four numbers, each of which is responsible for the breaking of a different type of symmetry (**Fig. 4a**). Using this reduced set, we perform a series of VFI simulations for symmetric and asymmetric embryo outline approximations. Six simulations are performed for each shape outline and an angular histogram for the lateral-anterior, ventro-lateral and posterior regions is generated (**Fig. 4b**). In addition to the approximate outlines, we include shapes with both their anterior and posterior features identical (a-p symmetric), but based on the original (experimental) outline (**Fig. 4c**). Finally, an average deviation score, as compared to the experimental tissue flows, is calculated (**Fig. 4d**). In all cases with a-p symmetric outlines no preferred pole for the dorsal movement was observed when considering the averaged simulations. Shapes that were a-p asymmetric, however, showed a clear preference for dorsal movement of posterior tissue (in all cases, *n* = 6). We conclude that the break of a-p symmetry (non-zero value of the 4^th^ shape parameter in **Fig. 4a**) of the egg, coupled to the positioning and geometry of the VFI-region, are responsible for differential tissue flow.

In addition, and related to the results above, the simulated tissue flow pattern qualitatively predicts the anterior-posterior anisotropy of the tissue of the VFI, which has been observed experimentally. Using SiMView recordings of fluorescently membrane-labeled whole-embryos (**Fig. 5a,b**), the data is segmented to obtain three-dimensional cell outlines (**Fig. 5c**). In a cross-section of the embryo (cut through and orthogonal to the middle of the a-p axis), we observe good agreement of changes in cell shapes (experimental) vs. simulated tissue deformation (**Fig. 5c-e**). Anisotropy with respect to the a-p axis is calculated for all segmented cells (experimental) and from the strain field (simulated) corresponding to changes within the first 10 minutes of onset of gastrulation, and mapped to the embryo outline, showing good agreement (**Fig. 5f,g**). It is important to note that, although the outcome of the simulation is anisotropic deformation, the simulation correctly used isotropic contraction of the cytoskeleton, by enforcing an isotropic preferred curvature change. This is in accord with observed isotropic contraction of the actomyosin meshwork determined experimentally (45). In the context of mechanical strain energy, this result can be explained as follows: due to the (isotropic) contractile forces of the actin-myosin meshwork, epithelial tissue surrounding the mesoderm primordium region is forced to extend towards the forming VFI. If this tissue is near the poles, then it has to change its curvature significantly (departing from its high preferred curvature). This comes at a high energy cost. It is easier (lower energy) for tissue to be “pulled” from the lateral parts of the embryo towards the furrow region, because only a relatively small change in curvature is associated with its extension towards the ventral side. This result suggests plasticity in the epithelium that sets in at a time prior to the onset of VFI and in which the egg serves as a scaffold, and indicates a direct mechanical influence that morphology of the egg potentially exerts on fly embryo development.

**Figure 5.**
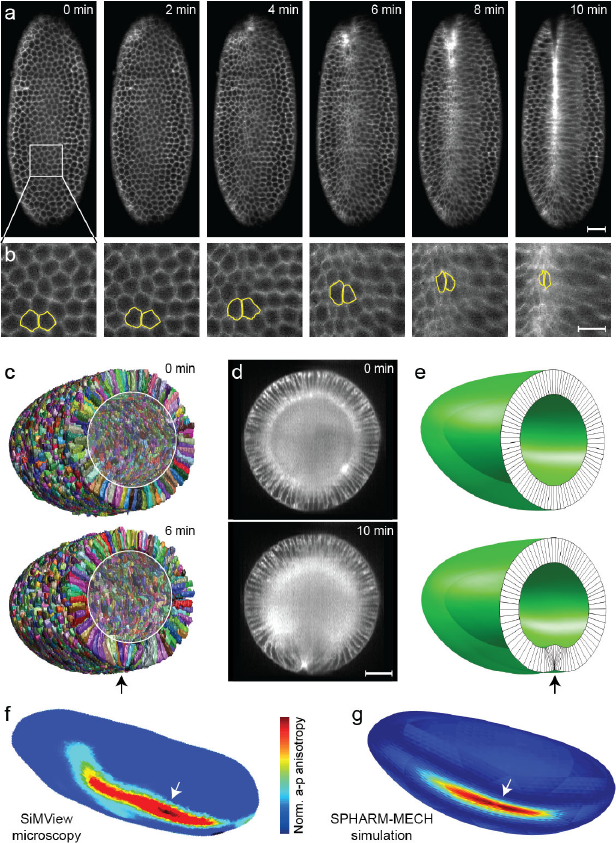
Comparison of experimentally observed and simulated tissue anisotropy. (**a**) Time sequence of Spider-GFP SiMView light-sheet microscopy images. Whole-embryo three-dimensional stacks were recorded (**Supplementary Video 1**). Only one plane, which approximately cuts through the ventral furrow of each stack, is shown, with the anterior pole at the top. (**b**) Enlarged image of the region marked in (a) of the same time sequence, following the outlines of two neighboring cells (yellow outlines), and showing the development of anisotropy as the ventral furrow invagination forms. (**c**) Segmentation results for time points 0 min and 6 min. Black arrow: beginning of furrow formation. (**d**) SiMView light-sheet microscopy images of Spider-GFP (membrane label) embryos. Approximately the middle planes of the three-dimensional stacks at time 0 min and 10 min are shown. (**e**) Corresponding simulation cut-through surfaces when the preferred local mean curvature of the mid-plane surface at the active region is increased from -0.5 (top) to 1.5 (bottom). Lines are shown as visual guides to the simulated deformation and should not be understood as cell boundaries. Black arrow: beginning of furrow formation. (**f**) Whole blastoderm morphology in perspective view with color code equal to a-p anisotropy (normalized). White arrow: beginning of furrow formation, corresponding to region indicated by black arrow in (c). (**g**) Same as (f) performed for the simulated morphology. White arrow: beginning of furrow formation, corresponding to region indicated by black arrow in (e). Scale bars, 20 μm (a), 10 μm (b), 50 μm (d).

## Application-Specific Methods

### Specimen preparation and live imaging

*Drosophila* live imaging experiments were performed with embryos homozygous for the membrane label Spider-GFP and the nuclear label His2Av-RFP. This line was constructed by combining stocks of w*; P{w[+mC]=His2Av-mRFP}; + (Bloomington *Drosophila* Stock Center, 23560) and w; +; Spider-GFP (29). Double labeled *Drosophila* embryos were dechorionated with 50% sodium hypochlorite solution (Sigma-Aldrich, 425044) and embedded in 1% low-melting temperature agarose (Lonza, SeaPlaque) in a 2 mm O.D. × 20 mm glass capillary (Hilgenberg GmbH) as has been previously described (42). After polymerization, the agarose cylinder was extruded just enough to expose the embryo outside of the glass capillary. The capillary holding the embryo was mounted vertically within the water-filled recording chamber of the SiMView light sheet microscope. Images were acquired at 2-minute intervals. Each time point comprises two-color z-stacks recorded from four orthogonal optical views for two different physical orientations (dorso-ventral and lateral), encompassing the entire volume of the embryo with an axial step size of 1.950 μm. For all data presented, the recording was terminated when the larva hatched and crawled out of the field of view (**Supplementary Movie 2**), after which the larva was transferred to a standard vial of fly food and raised to adulthood.

### Image processing and analysis

Prior to multiview fusion, raw SiMView recordings were corrected for insensitive pixels on the sCMOS detectors, using median and standard deviation filters. Multiview image fusion was performed with a custom Matlab processing pipeline, using rigid transformation for multiview stack registration followed by linear blending (42, 46).

Automatic segmentation of the membrane marker channel was performed with the watershed algorithm and subsequent agglomeration of oversegmented regions by persistence-based clustering (47). To remove false positives, the nuclei marker channel was segmented with the same methodology, and a one-to-one matching between nuclei and membrane segmented regions was obtained using the Jaccard distance and the Hungarian algorithm (48). Only objects that have a unique correspondence to the nuclei channel were assumed to correspond to cells and were used in the analysis. Cell segmentations were improved manually by an expert using itkSNAP. The centroids of the segmented cells were used to construct a point-cloud. Whole-embryo morphologies were obtained by interpolation of point-cloud data (**Supplementary Note 5**). Anisotropy with respect to the anterior-posterior axis was scored by approximating the cell outlines by ellipsoids, projecting the axes of these ellipsoids onto the plane parallel to the anterior-posterior axis, projecting the result onto the anterior-posterior axis and taking the ratio of largest to smallest projection.

## Summary

This work demonstrates whole-embryo mechanics modeling that is able to predict global changes in tissue flows and anisotropy based on local forces whose effect is expressed in terms of local preferred curvature. The approach was facilitated by a computational biomechanics framework that is data-driven and inherently three-dimensional, accommodates a large range of morphological, gene expression and material properties, is independent of data source, and unifies its analysis within a tissue mechanics context. The software used is made available online at https://github.com/khaledkhairy/SPHARM_Mech-Project (also see **Supplementary Note 9**). **Supplementary Videos 3-6** provide a brief introduction to the user interface. We envision that this approach can be applied to a wide spectrum of developmental biology model systems, and will facilitate the testing of effects of mechanical and genetic perturbation in a biomechanical context.

## Acknowledgements

We wish to thank Johannes Baumgart for his helpful comments on biomechanics modeling, Jonathan Howard for critical reading of the manuscript, Eric Wieschaus (HHMI) for providing the Spider-GFP flies, and Aleksandra Denisin for her contributions to implementing the SiMView imaging assay. This work was supported by the Howard Hughes Medical Institute.

---

**Supplementary Figure 1.**
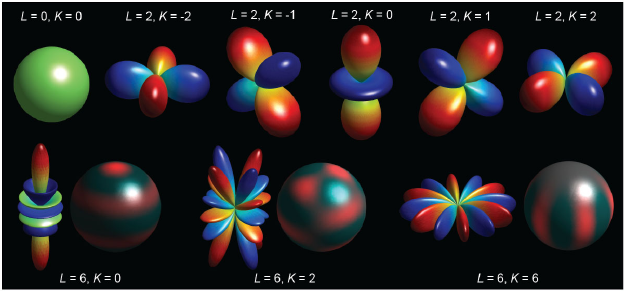
Spherical harmonics basis functions. Examples of spherical harmonics basis functions (**Supplementary Note 1**). These functions are well known as the angular portion of a set of solutions to Laplace’s equation. Color code: blue negative to red positive values. In the bottom row a perspective view of the corresponding mapping of function values to the unit sphere is shown.

**Supplementary Figure 2.**
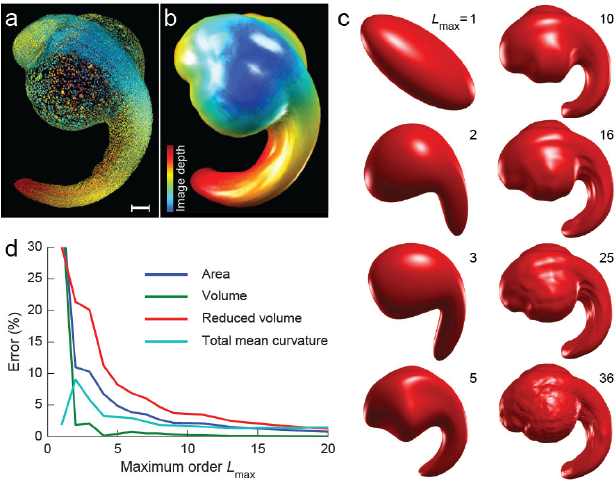
SPHARM representation of a zebrafish embryo. (**a**) Maximum-intensity projection of a SiMView image stack showing a 22 hours post fertilization zebrafish embryo. Color indicates depth into the image. (**b**) SPHARM-MECH reconstructed surface of (a) with maximum order *L*_max_ = 20. (**c**) Reconstructed surfaces using increasing *L*_max_. (**d**) Convergence of the spherical harmonics series (fidelity to the data): plot of percent error of surface properties of original surface mesh reconstruction *vs.* maximum order *L*_max_ used in fitting spherical harmonics coefficients. Scale bar, 100 μm.

**Supplementary Figure 3.**
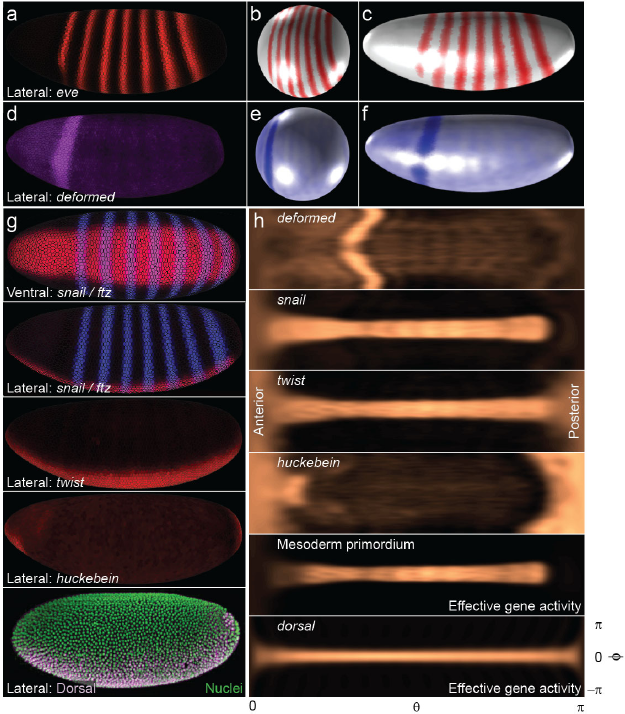
Mapping of gene expression patterns and determination of mesoderm primordium and active regions. (**a**) Expression pattern of *eve.* (**b**) Mapping of *eve* to the unit sphere. (**c**) SPHARM-MECH representation of *eve* shown on the full blastoderm morphology. (**d-f**) same as a-c but for *deformed* (*dfd*) which is used to define the region of activity for the cephalic furrow formation. (**g**) From top to bottom: ventral and lateral views of *ftz* and *snail* patterns, lateral view *twist*, lateral view *huckebein* and lateral view *dorsal.* Data for eve, *dfd, snail, twist* and *huckebein* is obtained from the Berkeley Virtual Embryo Project and displayed using PointCloudXplore Light. (**h**) Two-dimensional angular plots of the spherical harmonics representation of dfd, *snail, twist* and *huckebein* (top four panels), the interaction map of *snail, twist* and *huckebein* (**Supplementary Note 8**), and (bottom most) the pattern of *dorsal.*

**Supplementary Figure 4.**
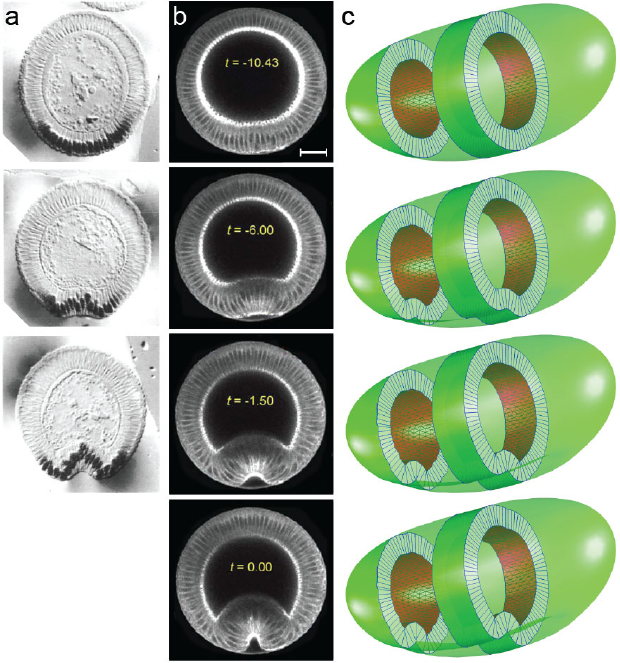
Ventral furrow invagination images compared to SPHARM-MECH simulation results. (**a**) *D. melanogaster* embryo electron microscopy images (adapted from Fig. 2 in Leptin and Grunewaldt (36)), and (**b**) multi-photon microscopy images of transgenic Sqh-GFP embryos (adapted from Fig. 6 in Conte *et al.* (17)). (**c**) SPHARM-MECH simulation of VFI showing a sequence of gradually increasing preferred curvatures when using the *dorsal*-based activity map of **Fig. 2a**. Scale bar, 20 μm.

**Supplementary Figure 5.**
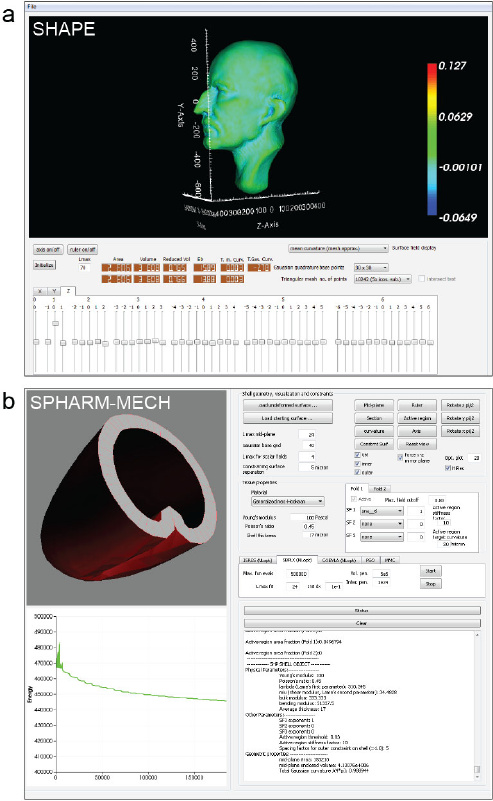
Screenshots of SHAPE and SPHARM-MECH. (**a**) **SHAPE** (**S**pherical **HA**rmonic **P**arameterization **E**xplorer) is a MacOSX 64bit compiled program with a graphical user interface (GUI) that allows the user to view and modify SPHARM shapes that have been saved in the “.shp3” format. “.shp3” stores morphology and scalar fields such as gene expression patterns in the form of spherical harmonics coefficients in a text file. The program also demonstrates calculation of surface properties with Gaussian quadrature in comparison to usage of the triangular mesh. SHAPE is based on the shape_tools library and uses in addition QT and VTK (Visualization Toolkit) C++ libraries. (**b**) **SPHARM-MECH** (SPHARM-Mechanics) is a MacOSX 64bit compiled program with a GUI that allows the user to import shapes in the “.shp3” format, configure a tissue shell mechanics calculation and execute numerical optimization. The result is a lowest mechanical energy shape prediction. The screenshot shows a predicted VFI morphology at the end of an optimization using the SubPlex algorithm. SPHARM-MECH is based on the shape_tools library.

